# Highly multiplexed spectral FLIM via physics informed data analysis

**DOI:** 10.1101/2025.08.04.668462

**Authors:** Mohamadreza Fazel, Reza Hoseini, Ayush Saurabh, Lance W. Q. Xu, Lorenzo Scipioni, Giulia Tedeschi, Enrico Gratton, Michelle A. Digman, Steve Pressé

## Abstract

Spectral fluorescence lifetime imaging (S-FLIM) allows for the simultaneous deconvolution of signal from multiple fluorophore species by leveraging both spectral and lifetime information. However, existing analyses still face multiple difficulties in decoding information collected from typical S-FLIM experiments. These include: using information from pre-calibrated spectra in environments that may differ from the cellular context in which S-FLIM experiments are performed; limitations in the ability to deconvolute species due to overlapping spectra; high photon budget requirements, typically about a hundred photons per pixel per species. Yet information on the spectra themselves are already encoded in the data and do not require pre-calibration. What is more, efficient photon-by-photon analyses are possible reducing both the required photon budget and making it possible to use larger budgets in order to discriminate small differences in spectra to resolve spatially co-localized fluorophore species. To achieve this, we propose a Bayesian S-FLIM framework capable of simultaneously learning spectra and lifetimes photon-by-photon ultimately using limited photon counts and being highly data efficient. We demonstrate the proposed framework using a range of synthetic and experimental data and show that it can deconvolve up to 9 species with heavily overlapped spectra.

## Introduction

Fluorescence microscopy techniques leverage multiple fluorophore properties to monitor subcellular structures and processes^1–1^. Among all these techniques, fluorescence lifetime imaging (FLIM)^11-14^ and spectral imaging^2,15–18^ have been extensively employed to study sub-cellular environments. However, these techniques are limited in multiplexed imaging due to their restriction in deconvolving a large number of fluorophore species based on either distinct fluorophore lifetimes or spectra. As such, spectral-FLIM (S-FLIM) integrates lifetime with spectral properties to differentiate a large number of fluorophore species and achieve highly multiplexed imaging^4 19 24^. For instance, S-FLIM has been exploited for simultaneous study of multiple different proteins, *e*.*g*., glucose transporter 1 (GluT1), and glial fibrillary acidic protein (GFAP), distribution within nerve cells to learn about their spatial organization, relative localization, and complex interactions in highly heterogeneous specimens such as the brain^22^. Additionally, spectral FLIM differentiates endogenous autofluorescent species for metabolic imaging based on their spectral decays, enabling detailed analysis of organelle interactions and biochemical processes across multiple cellular compartments simultaneously^15,2^. Another example includes the study of breast cancer via abundance of key endogenous components, such as nicotinamide adenine dinucleotide (NAD) and its different forms, as well as flavin adenine dinucleotide (FAD) involved in the metabolism of human breast cell lines^12^.

In typical S-FLIM experiments, the sample is raster scanned, spot-by-spot, with a pulsed laser. At each spot, fluorophores can be excited during laser pulses leading to photon emissions which are, in turn, passed through a diffraction grating to disperse the light into multiple spectral bands (channels) with 10-20 nm widths; see Fig. 1. Subsequently, a single photon detector records the photon counts recorded for each spectral band, and the corresponding photon arrival times with respect to the laser pulse centers (microtimes); see Fig. 1. More concretely, the recorded photon arrival times typically differ from the exact time that a fluorophore spends in the excited state due to the finite laser pulse width and detector delay. These effects together are termed the instrument response function (IRF) and contribute to the noise in the set of collected arrival times ^13^. Moreover, photon counts for each band deviate from the expected counts, *i*.*e*., shot noise. As such, the acquired data set, while inherently stochastic, contains a set of detected photons, each stamped with the corresponding spatial dimensions (*x, y*), a spectral band, and an arrival time.

**Figure 1:**
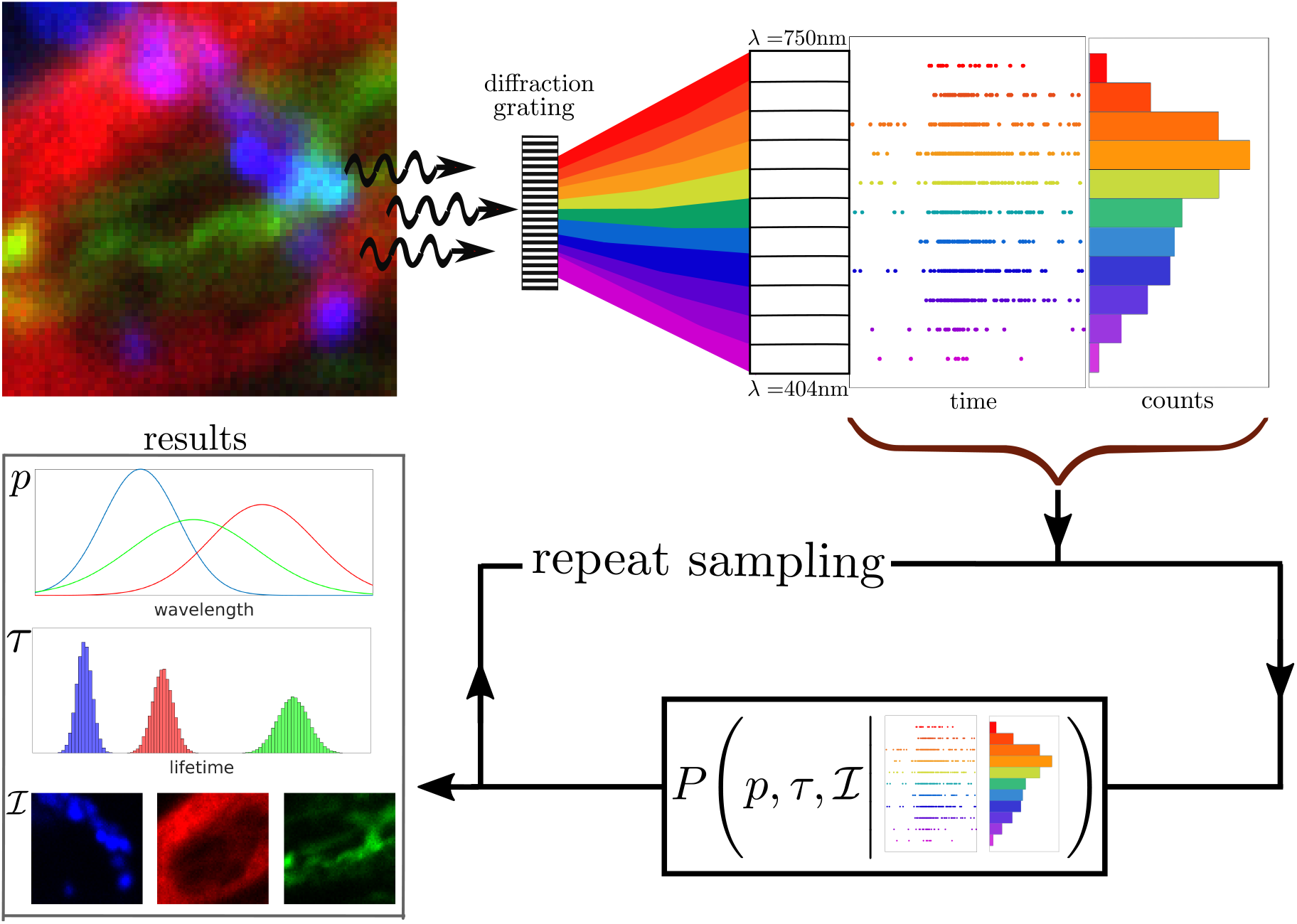
Conceptual workflow of a spectral-FLIM experiment and our framework. Starting from the top left, the sample is scanned using a confocal setup leading to photon emissions from fluorophore species (here shown in red, green, and blue) staining different structures. The emitted photons pass through a diffraction grating and undergo dispersion based on their wavelengths. The photons are then detected within one of the spectral channels based on the dispersion angle. The photon arrival times and photon counts from each channel are recorded and used to construct the central object of interest here: the spectral FLIM posterior. This posterior can then be iteratively sampled to infer spectra, lifetimes, and fluorophore distribution maps, respectively, associated to each fluorophore species.

The set of acquired data from all species combined (photon arrival times and spectra) can be deconvolved to learn the spectra and lifetimes of each fluorophore species, and infer the underlying spatial fluorophore distribution maps. For instance, a few FLIM analyses methods have been adapted for S-FLIM data analyses including: phasor^3,4 17 25^; and linear and non-linear mixture models^22,26^. However, these approaches face several challenges, including: 1) their ability to handle many overlapping species due to lack of rigorous treatment of uncertainty. For instance, phasor S-FLIM can handle up to three spatially overlapping species^4^; 2) requirement to pre-calibrate or assume specific forms, *e*.*g*., Gaussian^17^, for the spectra; 3) their limitation in deconvolving fluorophore species with high spatial and spectral overlap and similar lifetimes; 4) high photon count requirement due to inaccurate modeling of noises involved in the problem, *e*.*g*., detector and photon shot noises.

To overcome the challenges above, we propose a novel S-FLIM framework jointly modeling emission spectra and lifetime to facilitate highly multiplexed imaging. To achieve this, our method takes traces of photon arrival times and photon counts per spectral band for each spot to simultaneously deduce species lifetimes, spectra, and the corresponding fluorophore distribution maps tied to the local concentration of species at each spot.

Moreover, for accurate characterization of the species’ spectra, lifetimes, and the rigorous propagation of uncertainty from sources such as IRF and Poisson noise, we adopt a Bayesian framework. More specifically, we employ Dirichlet priors^13,27,28^ to deal with inherently noisy spectral data. Moreover, working within the Bayesian framework, our method is capable of operating under photon starved conditions or regions of images with naturally low counts, common for biological samples, and dealing with highly spectrally and spatially overlapped fluorophore species.

In what follows, we put forward the mathematical description of our Bayesian S-FLIM framework. We validate our method through simulations assuming multiple fluorophore species intentionally selected as being un-analyzable using other tools and experiments on microtubules stained with ViaFluor 488, mitochondria stained with MitoTracker Orange, and lysosomes stained with LysoTracker Red given low photon count data, demonstrating superior resolution of a greater number of fluorophore species for highly multiplexed biological imaging with heavily overlapped spectra.

## Results

Our S-FLIM framework’s main goal is to simultaneously learn lifetimes, *τ*_1:*M*_ and spectra, ***π***_1:*M*_, of *M* fluorophore species present within the data and the corresponding fluorophore distribution maps, 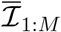. To do so, we need to operate within a Bayesian framework and work with a posterior, which assumes a non-analytic form; see Section. We use a Monte Carlo procedure to numerically sample the posterior; see Section. Therefore, the results presented in this section are either histograms of these drawn samples or the corresponding posterior median.

Here, we use both synthetic and experimental data to benchmark our S-FLIM framework. First, we employ synthetic data to show that relying on only lifetime information is not sufficient to deconvolve 4 fluorophore species (see Fig. S1). We next employ synthetic data to demonstrate our S-FLIM framework’s ability to: 1) deconvolve overlapping fluorophore maps (see Fig. 2); 2) learn maps for heavily overlapping spectra (see Figs. 2 and S2); 3) infer lifetimes below the IRF and with sub-nanosecond differences (see Figs. 2 and S2); 4) deal with highly multiplexed data containing as many as 9 different fluorophore species (see Fig. S2); and 5) surpassing current S-FLIM data analysis methods^4 23^ in learning lifetimes and spectra using as few as 5000 photons (see Fig. 3 ; Finally, we employ experimental data to assess the robustness of our S-FLIM framework in deducing different lifetimes and overlapping spectra with intricate fluorophore maps, (see Figs. 4 and S3).

**Figure 2:**
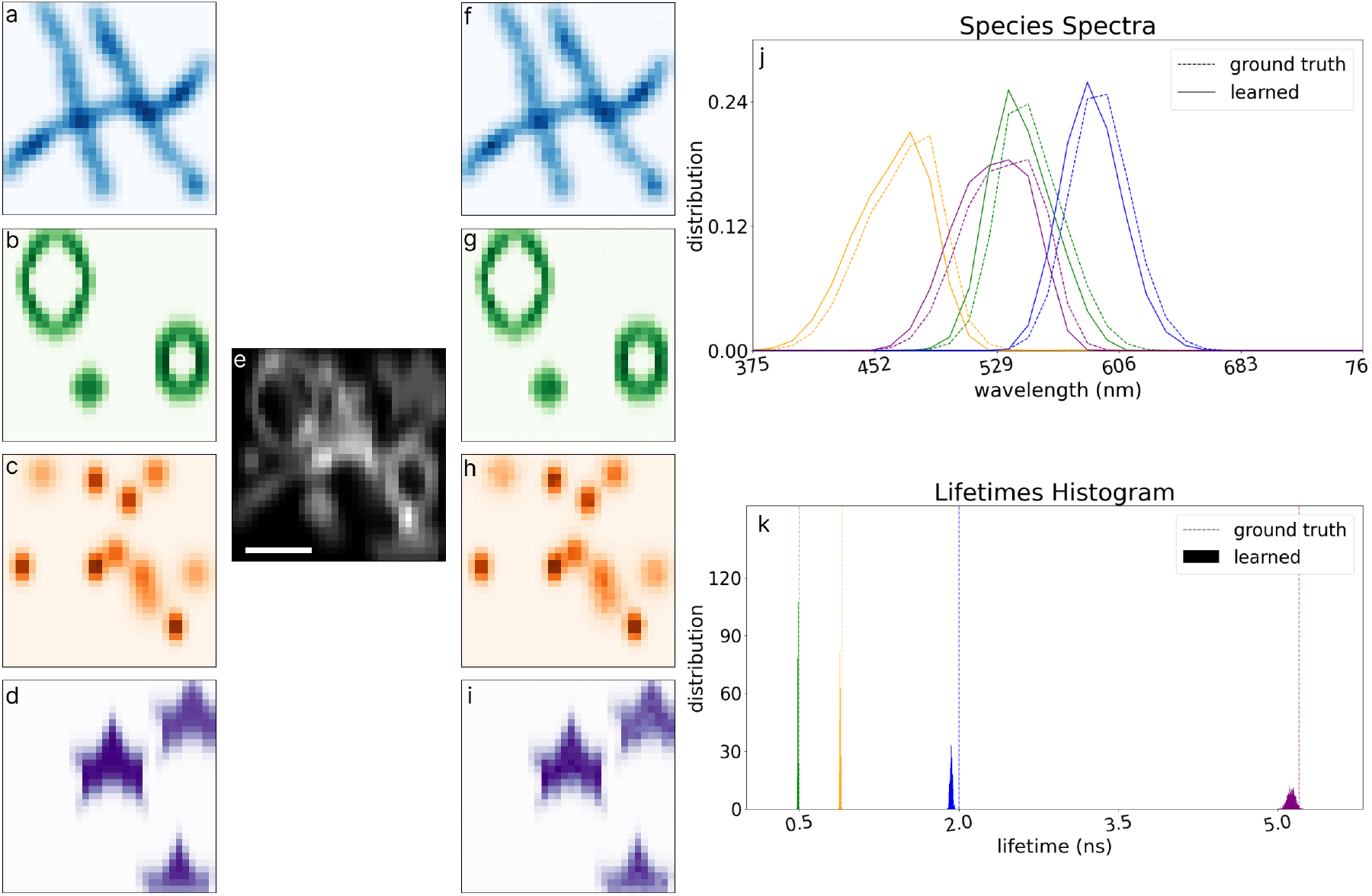
Fluorophore distribution deconvolution using both lifetime and spectral information. (a-d) 4 fluorophore distribution maps synthesized over an area of 32 *×* 32 pixels; (e) simulated data using the fluorophore distibutions in panels a-d; (f-i) deconvolved fluorophore distributions maps using both lifetime and spectral information; (j) mean of the sampled spectra; (k) histogram of the lifetime samples. Scale bar is 2 *μm*.

**Figure 3:**
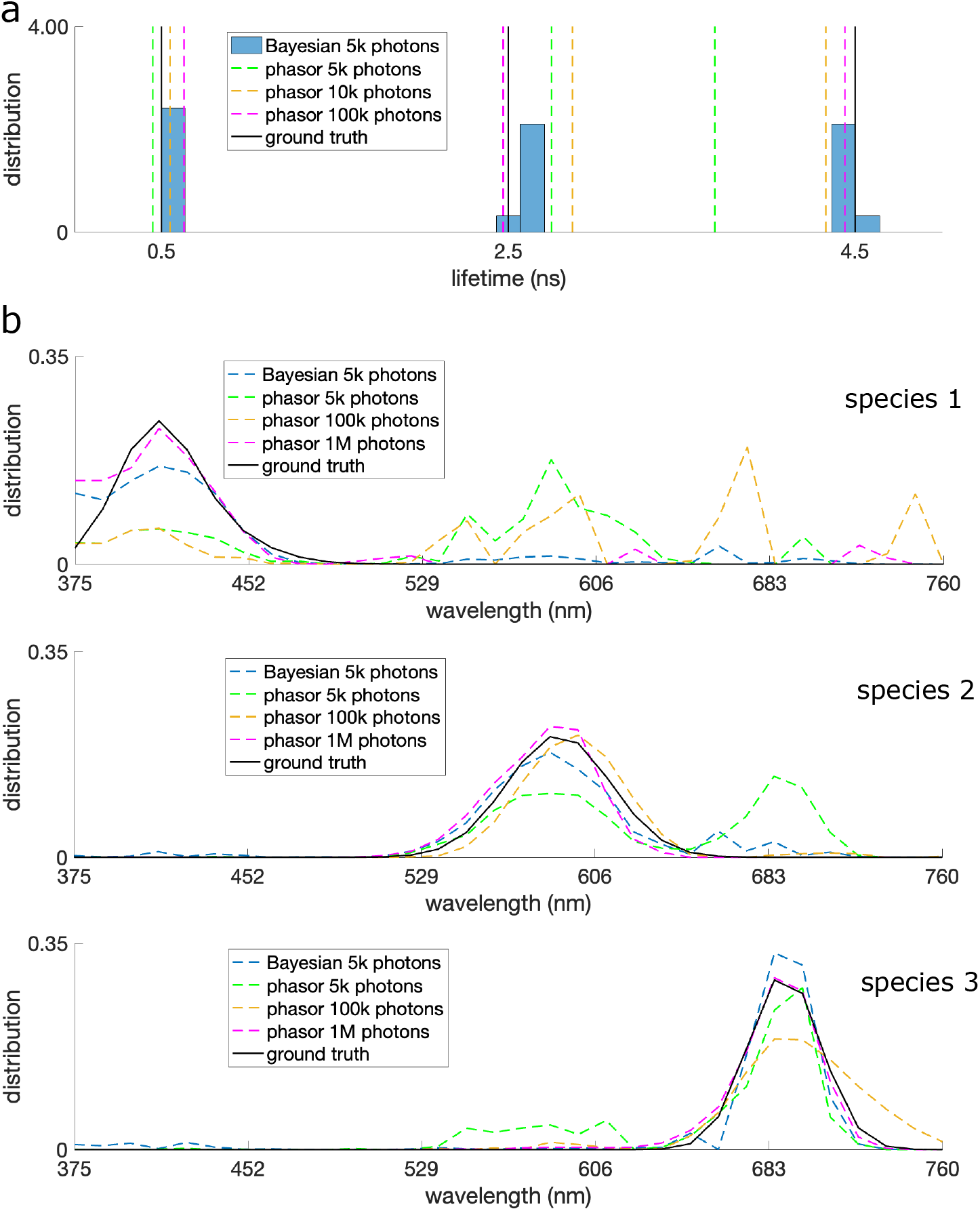
Comparing phasor and Bayesian S-FLIM frameworks using different photon counts. (a) Histograms show the sampled lifetimes from our Bayesian method using 5k photons with the green, yellow, and red dashed lines coinciding with lifetimes found by the phasor method using 5k, 10k, and 100k photons, respectively. The black lines designated the ground truth lifetimes. (b) The black lines in the top, middle and bottom plot show ground truth spectra for the first, second, and third species corresponding to lifetimes of 0.5 ns, 2.5 ns and 4.5 ns. The blue dashed lines represent the median of sampled spectra by our Bayesian method. The green, yellow, and red dashed lines show the spectra recovered by the phasor method using 5k, 10k, and 100k photons.

**Figure 4:**
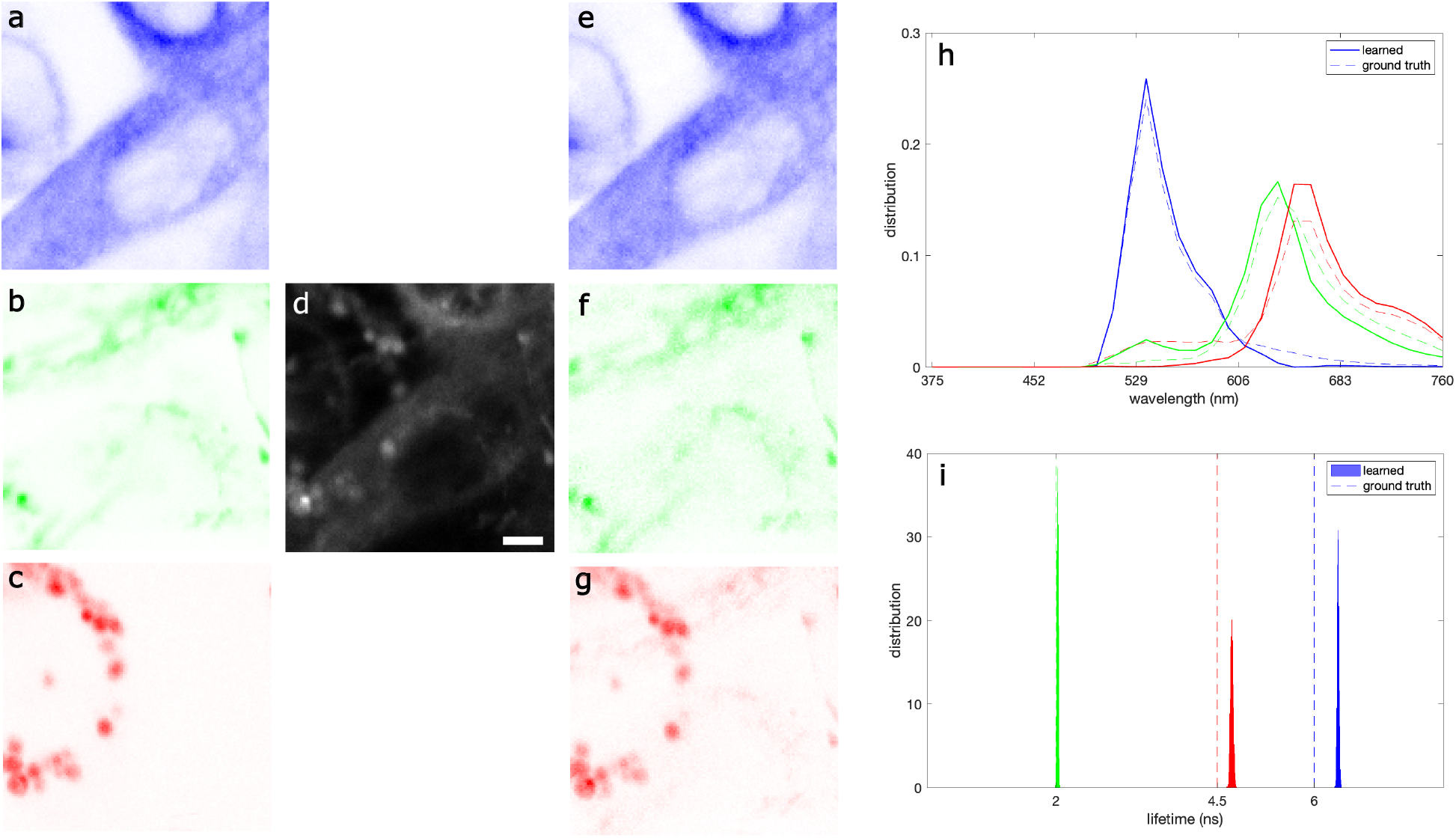
To help validate our method on realistic fluorophores distribution maps, we consider experimental S-FLIM data by mixing three single-species S-FLIM data sets (panels a-c) into one. That is, we first analyze three data sets each containing a single species to produce the “ground truth” lifetimes and spectra shown in panels (h) and (i). (a) single-species data of microtubules stained with viaFluor 488 with a lifetime of 6 ns–shown in blue; (b) single-species data of mitochondria stained with MitoTracker Orange with a lifetime of 2 ns–shown in green; (c) single-species data of lysosomes stained with LysoTracker Red with a lifetime of 4.5 ns–shown in red; (d) the three data sets in (a-c) are combined to generate a pseudo data with 3 species; (e-g) deconvolved fluorophore distribution maps using our S-FLIM framework; (h) mean of the sampled spectra; and (i) histograms of the sampled lifetimes. Scael bar is 2 *μm*.

### Robustness using simulated data

Here, we begin by demonstrating the advantage of integration of spectral information into the analysis. To do so, we use four synthetic fluorophore maps (ground truth fluorophore maps) over an area of 32*×*32 pixels (see Figs. 2 and S1) with lifetimes of 0.4 ns, 0.9 ns, 2 ns and 5 ns to simulate S-FLIM data as described in Section ; see Figs. 2(e) and S1(e). Using the synthesized maps, we simulated 272K photons in total with 92K, 60K 48K and 72K photons from maps shown in Fig. 2a-d. We process the resulting dataset by both ignoring (Fig. S1) and considering (Fig. 2) the spectral information and compare the results. Indeed, panels Fig. S1(f-i) illustrate the resulting fluorophore maps demonstrating failure to recover ground truth fluorophore maps in panels (a-d) when discarding the spectral information. The histogram of the sampled lifetimes is also depicted in Fig. S1j displaying a wide range of values instead of sharp distributions centered around the known ground truth values.

Next, we processed the same simulated dataset while incorporating the spectral information in addition to lifetime information. The resulting fluorophore maps are represented in Fig. 2(f-g). For quantitative comparison of these resulting fluorophore maps and the ground truth fluorophore maps in Fig. 2(a-d), we calculated the resulting errors by finding their differences given by Δℐ *×* 100*/*max(ℐ_true_), where ℐ_true_ indicates the ground truth fluorophore map. The difference maps are shown in Fig. S3(a) with average differences of *≈* 5%, *≈* 2%, *≈* 3% and *≈* 6% for the fluorophore maps shown in blue, green, red, and violet in Fig. 2. Fig. 2(j) depicts the ground truth and learned spectra corresponding to the fluorophore maps. Here, the average error in learning the spectra is less than 3%. The sampled lifetimes histograms are shown in Fig. 2(k). Our S-FLIM framework has learned all four lifetimes including the lifetime below the IRF (0.2 ns) with an average error of *≈* 2.5% when incorporating spectral information into the analysis.

Next, we employ synthetic data to benchmark our Bayesian S-FLIM’s sensitivity to photon numbers used in the analysis. To this end, we simulated data sets with 5k, 100k, and 1M photons over 10 pixels assuming 3 fluorophores species with lifetimes of 0.5 ns, 2.5 ns and 4.5 ns (see Fig. 3. Our Bayesian S-FLIM framework learns the correct lifetimes and the corresponding spectra using as little as 5k photons (see Fig. 3. In contrast, the phasor S-FLIM method^4^ requires more than 100k photons under the same conditions to learn the correct spectra (see Fig. 3b).

Finally, we used simulated data to demonstrate the capability of our S-FLIM framework to deal with the challenging case of highly multiplexed data containing 9 simultaneous fluorophore species. To do so, we simulated S-FLIM data (see Section) with 9 fluorophore species with approximately 60K photons from each species assuming heavily overlapped spectra with variable shapes, fluorophore distribution maps over an area of 32*×*32 pixels, lifetimes with sub-nanosecond differences, and an interpulse window of 14.9 ns; see Fig. S2. Fig. S2(a) depicts the learned and ground truth spectra with an average difference of *≈* 4%. Fig. S2b shows the corresponding histograms of sampled lifetimes. For lifetimes smaller than 2 ns the ground truth values fall within the range of the corresponding histograms. In contrast, for lifetimes larger than 2 ns, while the histogram peaks are close to the ground truth values with an average error of *≈* 8%, the ground truth lifetimes do not fall within the range of the corresponding histograms. This issue arises because of a flatter posterior for larger lifetimes due to larger number of photons from those species being detected after the pulse inducing the excitation making inference of the larger lifetime much more challenging ^13^; see Eq. 12.

### Robustness using experimental data

In this section, we demonstrate the capability of our S-FLIM framework to deal with intricate fluorophore distribution maps within sub-cellular environments. We start by evaluating our S-FLIM framework in deconvolving fluorophore distribution maps and learning the corresponding lifetimes and spectra. To do so, we use three experimental data sets each acquired over an area of 128*×*128 pixels and labeled by a single fluorophore species at a time including: ViaFluor 488 staining microtubules (shown in blue in Fig. 4); and MitoTracker Orange staining Mitochondria (shown in green in Fig. 4); and LysoTracker Red staining lysosomes (shown in red in Fig. 4); see Section. As these data sets were acquired using the same imaging setup under the same experimental conditions, as a control we mixed them to obtain a hybrid data set containing three fluorophore species; see Fig. 4(d). For this control, the individual spatial distributions of each fluorophore species is known. We then analyzed the hybrid S-FLIM data sets generated and compared the results with “ground truth”, *i*.*e*., the individual spatial distributions of each fluorophore species, as well as lifetimes and spectra obtained using data sets with individual fluorophore species. The resulting fluorophore distribution maps from our framework are shown in Fig. 4(e-g).

To quantify the robustness of our algorithm, we quantified the differences between the fluorophore maps obtained (Fig. 4(e-g)) and the ground truth fluorophore maps (Fig. 4(a-c)) as described for synthetic data in Fig. 2. The resulting difference maps are depicted in Fig. S4. The average errors for the fluorophore species shown in blue, green, and red are, respectively, *≈* 10%, *≈* 4% and *≈* 3%. The learned spectra are shown in Fig. 4(h) where the average difference of the learned and ground truth spectra are *≈* 14%. Fig. 4(i) depicts the sample lifetime histograms. The histogram of the two larger lifetimes deviates from the ground truth values by 0.25 ns and 0.4 ns for the species shown in red and blue. This is again because, for larger lifetimes, the posterior tends to become flatter due to larger numbers of photons being detected after pulses that follow the one that gave rise to their excitation. In fact, this hinders our ability to distinguish longer lifetimes with small differences on the order of *≈* 1 *ns*. However, with both spectral and lifetime information combined, even if larger lifetimes approaching the inter-pulse window can introduce complications, we still deduced spatial maps with low uncertainty.

## Methods

In this section, we first outline the data likelihood for Bayesian S-FLIM. We then discuss our inverse model framework and the data simulation procedure. For more details we refer back to the Supplementary Information.

### Model formulation

The input data to our S-FLIM analysis method is a set of stochastic arrival times 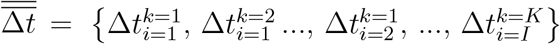; and a set of noisy photon counts 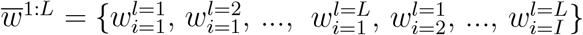 for *L* spectral bands, *I* pixels, and *K* photon arrival times in pixel *i*. Here, overbars represent the set of all possible values for either *i* or *k*. Our framework uses these input data to deduce the unknown parameters including lifetime values *τ*_*m*_ for species *m*, the corresponding expected photon count ℐ_*mi*_ in pixel *i* (the collection of all ℐ_*mi*_ across all pixels gives fluorophore distribution map for species *m*), and 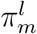 gives the area under the associated spectrum over spectral band *l*

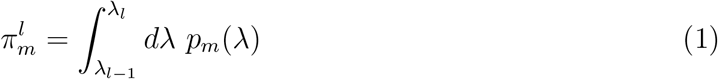

where *p*_*m*_(*λ*) is the normalized emission spectral (wavelength) distribution for *m*th species. To simplify the notation, we represent the set of 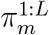 by ***π***_*m*_ here after.

The stochastic input data described above is probabilistically linked to the set of unknown parameters through the likelihood model

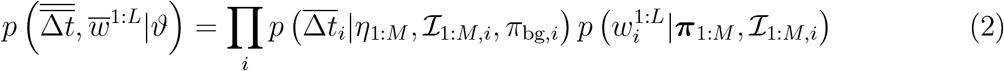

where *M* is the number of species, and *η*_*m*_ = 1*/τ*_*m*_. Here, we show the set of unknowns by 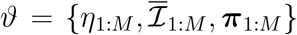, where the overbar indicates all the pixels. The two terms on the right hand side are

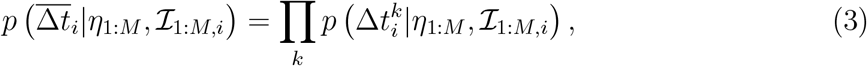

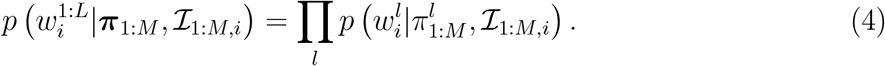

Here, 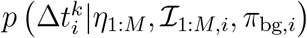 denotes the likelihood model for a single photon arrival time derived by considering a Gaussian IRF and exponential decay of the excited state while summing over all possible species and previous laser pulses that could give rise to the photon^13,14,29^

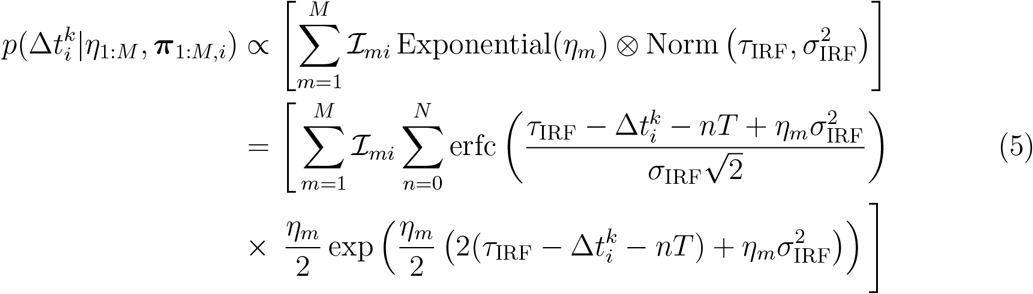

where *N, T, τ*_IRF_ and 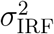 are, respectively, the maximum number of pulses over which a fluorophore may remain excited, laser interpulse time, offset and variance of the Gaussian IRF. Moreover, 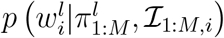 represents the likelihood of the photon count from spectral band *l* at pixel *i* given by a Poisson distribution

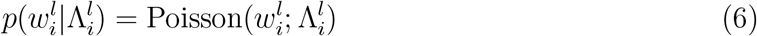

where the total expected photon count within the *i*th pixel and *l*th spectral band is given by

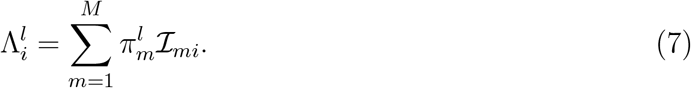

### Inverse model

In the last section, we derived the likelihood model for our spectral FLIM framework. In this section, we employ this likelihood in order to build the object of central interest within our Bayesian framework; namely the posterior. The posterior is proportional to the product of the likelihood and priors over the unknown parameters

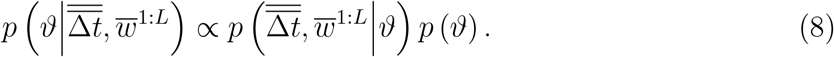

Here, we select a set of physically motivated priors. These include: Dirichlet priors, a generalization of the Beta distribution with more than two options ^13 28 30 32^, over ***π***_*m*_ to maintain the normalization of the emission spectrum *p*_*m*_(*λ*); Gamma priors over the inverse lifetimes *η*_1:*M*_ and photon counts ℐ_1:*M,i*_ to guarantee positive values

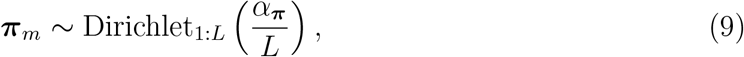

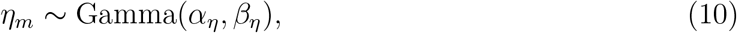

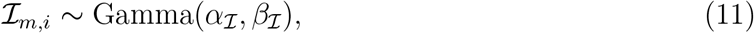

where *α*_***π***_, *α*_*η*_, *β*_*η*_, *α*_*ℐ*_ and *β*_*ℐ*_ are prior parameters.

The full posterior obtained does not attain an analytic standard form lending itself to direct sampling. Therefore, to draw inferences, we must numerically draw samples the posterior employing Markov Chain Monte Carlo techniques^10 14 28 33 35^. More specifically, we construct an overall Gibbs sampling scheme in order to sweep over all parameters at each iteration. This procedure is briefly outlined below and further detailed in the Supplementary Information:

1. inverse lifetimes *η*_1:*M*_ –these parameters are sampled using a Metropolis-Hasting (MH) scheme^36 37^.
2. spectral integrals, ***π***_*m*_–these parameters are also sampled using MH scheme as the conditional posterior does not attain a standard form.
3. photon counts, *ℐ*_*mi*_– these parameters are sampled again using the MH scheme.

Finally, the chain of drawn samples are used for further numerical analyses.

### Data simulation

In this section, we explain our synthetic data simulation procedure required in order to first demonstrate the method on synthetic data where ground truth is known. To generate synthetic photon arrival time traces at the appropriate wavelength, we first sample what species is excited. Next, we sample the excited state lifetime, Δ*t*_em,*k*_, from an exponential distribution and draw the wavelength from the given spectrum associated to the sampled species. Finally, we sample a stochastic IRF time, Δ*t*_IRF,*k*_, originating from the finite width of the laser pulse and detector delay. More specifically, we sample the fluorophore species resulting in the *k*th photon emission from a Categorical distribution, *i*.*e*., an extension to Bernoulli distribution with more than two options. Next, we sample the emitted photon wavelength and the excited state lifetime from the given associated spectrum and the exponential distribution with the corresponding lifetime, respectively. We then sample the IRF time from a normal distribution with mean and standard deviation of 2.5 ns and 0.5 ns similar to experimental data used in this study. Further, we used *T* = 12.8 ns for the interpulse time except when mentioned otherwise. Finally, for the case where the simulated photon arrival times exceed the interpulse time, *T*, we modify the arrival times by introducing the last term of the expression below

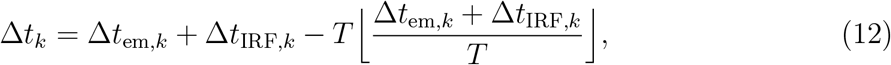

where ⌊ · ⌋ returns the integer portion of its argument.

This procedure is repeated for all pixels within the simulated data.

### Experimental Methods

All data were acquired with a custom-built microscope described in Ref. ^4^. Cells were excited with a 488-nm excitation light and a 500-nm long pass dichroic mirror (Semrock) through a Olympus PlanApo N*×*60/1.45 NA Oil objective. MDA-MB-231 cells were cultured in DMEM medium (DMEM, high glucose, with L-glutamine, GenClone), supplemented with 10% v/v FBS (heat inactivated FBS, GenClone) and 1% v/v penicillin/streptomycin solution 100*×* (10,000 units of penicillin and 10 mg/ml streptomycin in 0.85% saline solution, GenClone), in a 37 °C and 5% CO2 incubator and seeded in an eight-well chambered coverglass (Thermo Scientific, Nunc Lab-Tek), previously coated with 2 *μ*g/ml fibronectin in Dulbecco’s phosphate buffer solution (fibronectin human plasma 0.1% solution, Sigma-Aldrich; DPBS 1*×* without Ca, Mg, Phenol Red, GenClone).

The set of solutions used for calibration was prepared at a concentration of 7 *μ*M in PBS pH 7.4 and consist in Alexa 405 (Thermofisher Scientific), Fluorescein (Thermofisher Scientific) and ATTO490LS (ATTO-tec). Live cells were stained with 1:5,000 ViaFluor 488 (Biotium), 200 nM MitoTracker Orange CMTMRos (Thermofisher) and 1 *μ*M LysoTracker DND-99 Red (Thermofisher), incubated 30 min and washed twice before imaging. Dyes were diluted to have less than 0.1% DMSO in the medium.

## Discussion

Fluorescence S-FLIM microscopy integrates lifetime and spectral properties of fluorescent probes to achieve highly multiplexed imaging^4 20 24^. As such, this imaging technique is useful in simultaneously observing multiple sub-cellular processes and studying spatial distribution of macro-molecules across the cell. For instance, S-FLIM has been employed in spatial transcriptomics^2^ and to study the distribution of NAD and FAD within cells important in cancer cell metabolism.

Our S-FLIM framework offers a tool capable of dealing with highly multiplexed data containing many fluorophore species. In doing so, our Bayesian framework uses combined signal from all species (individual photon arrival times and photon counts per spectral channel) to simultaneously learn the associated lifetimes and spectra in order to infer fluorophore distribution maps. Moreover, our Bayesian framework addresses the existing challenges of current S-FLIM data analysis pipelines by relaxing the need to: assume Gaussian spectra^17^, dealing with overlapping spectra, spectral shifts in cellular environments^38^, and limitation in the number of fluorophore species.

We benchmarked our Bayesian S-FLIM framework using both synthetic and *in vivo* data. Using synthetic data, we showed that our method can learn spatial fluorophore maps even for heavily overlapping spectra and even in cases involving lifetimes below the IRF (see Fig. 2). We also demonstrated the ability of our code in dealing with highly multiplexed case by applying our method to a challenging S-FLIM data simulated assuming 9 fluorophore species with overlapping spectra (see Fig. S3). Finally, we used our method to analyze biological data sets containing heavily overlapped fluorophore distribution maps with intricate patterns (see Figs. 2 and 4). We showed that our method recovers the underlying fluorophore maps with a few percent error on average.

By nature, sequential Monte Carlo sampling is computationally costly. As such, we implemented our S-FLIM framework on graphic processor units (GPUs) where each processor calculated likelihood associated to a single pixel to speed up the likelihood calculations. To analyze data sets used in this work, we used a desktop with i9-14900KF processor, 64GB RAM and an RTX-4090 24GB GPU. Running the algorithm on the GPU, it took approximately 8 hours to process the experimental data set in Fig. 4. Further, for simulated data in Fig. 2, we had smaller computational times in order of less than 1 hour.

The framework developed in this work can be extended to learn IRF parameters, namely offset and standard deviation, by placing (most likely gamma) priors on these parameters and expanding our Gibbs sampling procedure. Furthermore, while here we only considered visible light, our method is capable of dealing with any spectral range. While we assumed known number of fluorophore species, our framework can be extended to the Bayesian nonparametric limit, at computational cost, to infer the number of species applicable to cases where these are unknown.

## Contributions

MF developed the mathematical model and algorithm. RH wrote the codes and performed the analyses with helps from MF, AS and LWQX. MF wrote the manuscript and the other authors contributed edits. LS, GT, MD and EG provided experimental data and useful suggestions along the way. SP supervised and oversaw the entire project.

## acknowledgment

S.P. acknowledges the support from NIH Grants R01GM134426, R01GM130745, and NIH MIRA R35GM148237. Image and data acquisition were made possible through access to the Laboratory for Fluorescence Dynamics, a shared resource center supported by the National Institutes of Health (grant no. P41GM103540 to L.S., A.V., and E.G.). This study was supported in part by funds from the National Science Foundation (grant nos. DMS1763272 and 1847005 to M.A.D.) and a grant from the Simons Foundation (594598 QN to M.A.D.).

## Code Availability

The software package is available on Github: Bayesian spectral-FLIM software package.

## Supplementary Information

**Fig. S1:**
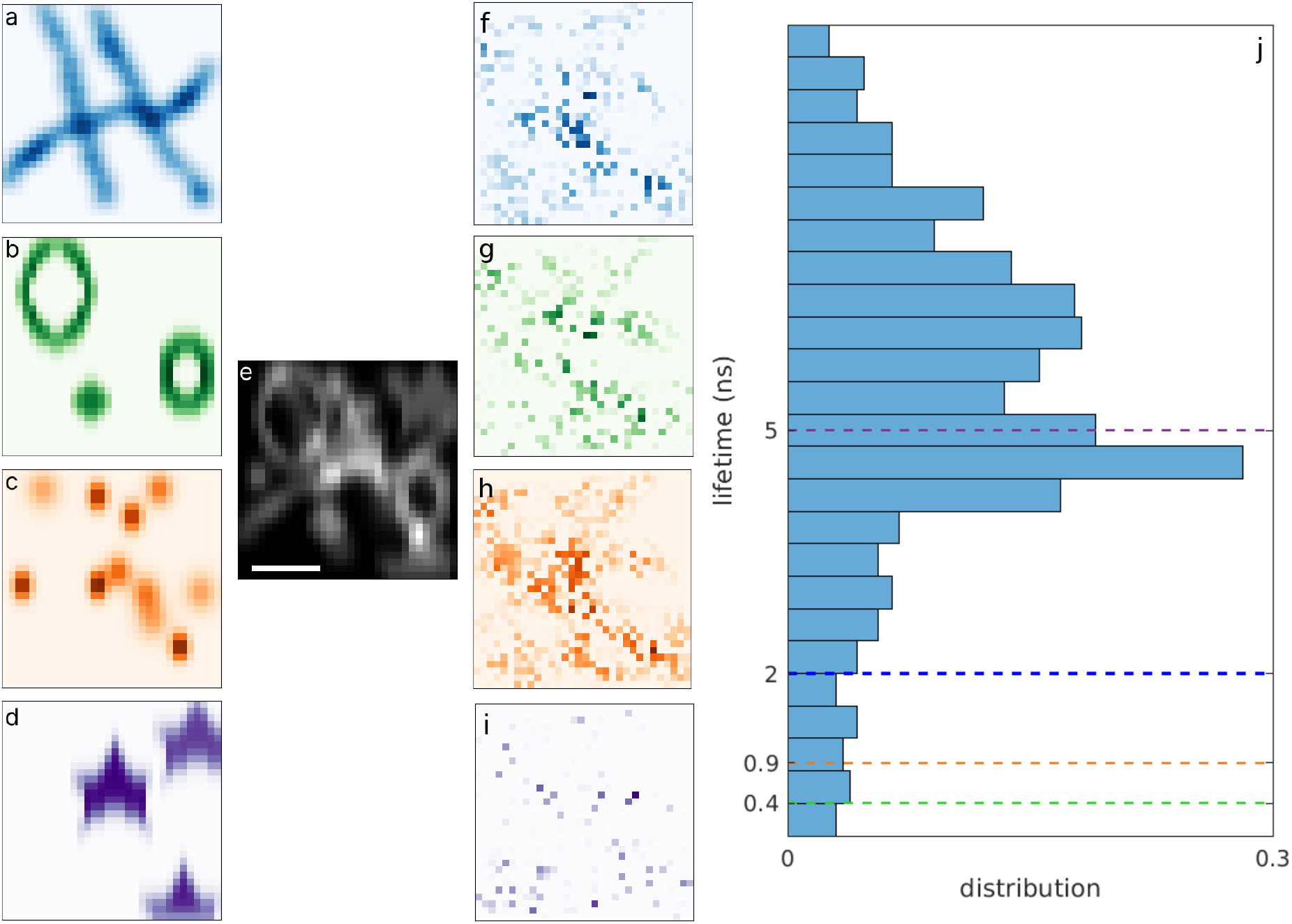
The failure of fluorophore distribution deconvolution using only lifetime information. (a-d) Four fluorophore distribution maps synthesized over an area of 32 *×* 32 pixels. The simulated data is the same as Fig. 2. (e) simulated data using the fluorophore distributions in panels a-d; (f-i) deconvolved fluorophore distribution maps using only lifetime information; (j) histogram of the lifetime samples. Dashed lines show the ground truths. Scale bar is 2 *μm*.

**Fig. S2:**
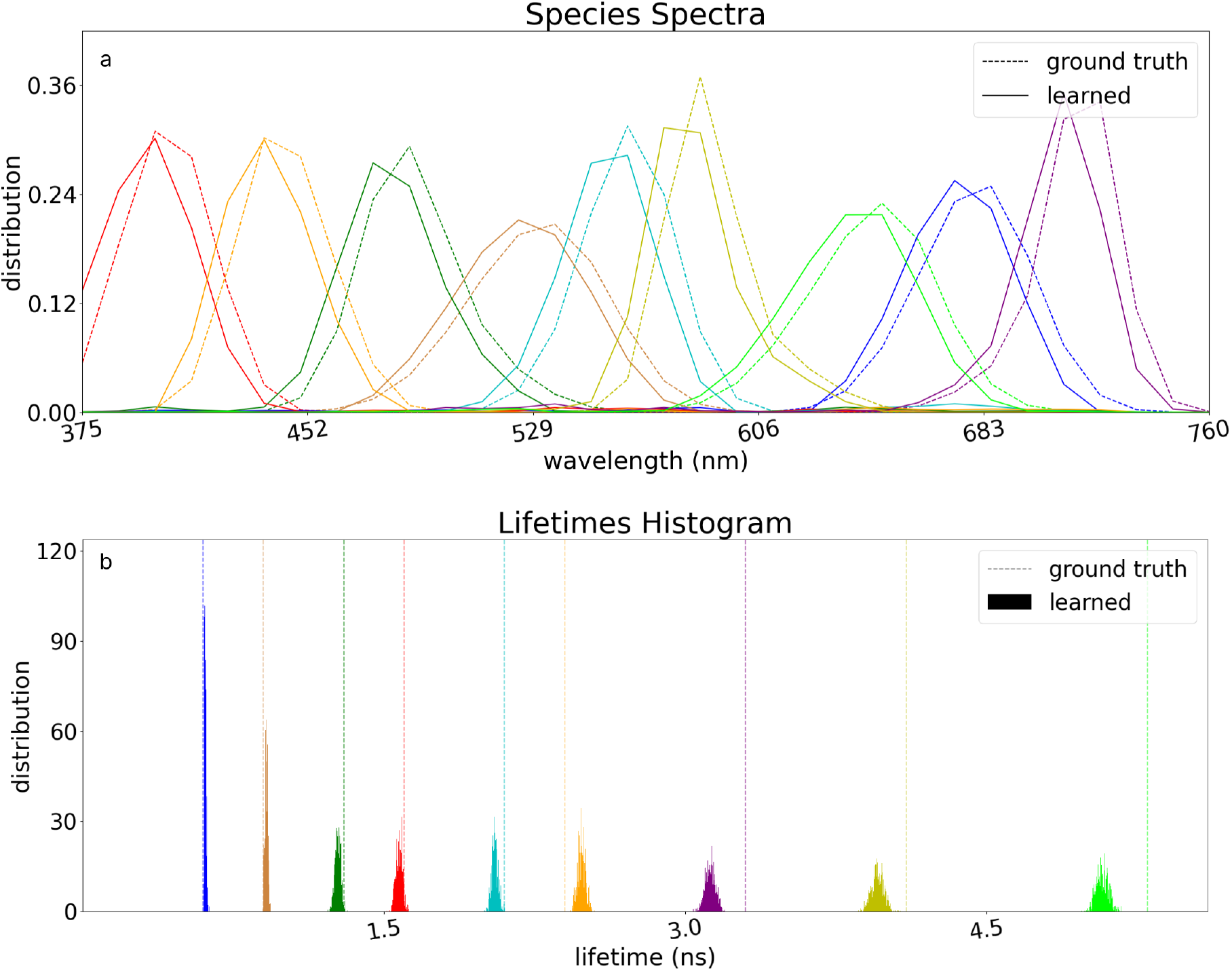
Learning lifetimes and spectra from highly multiplexed data containing 9 species. (a) Mean of the sampled spectra; (b) histogram of the sampled lifetimes. The total simulated photons are 540K with approximately 60K photons from each species over an area of 32 *×* 32 pixels.

**Fig. S3:**
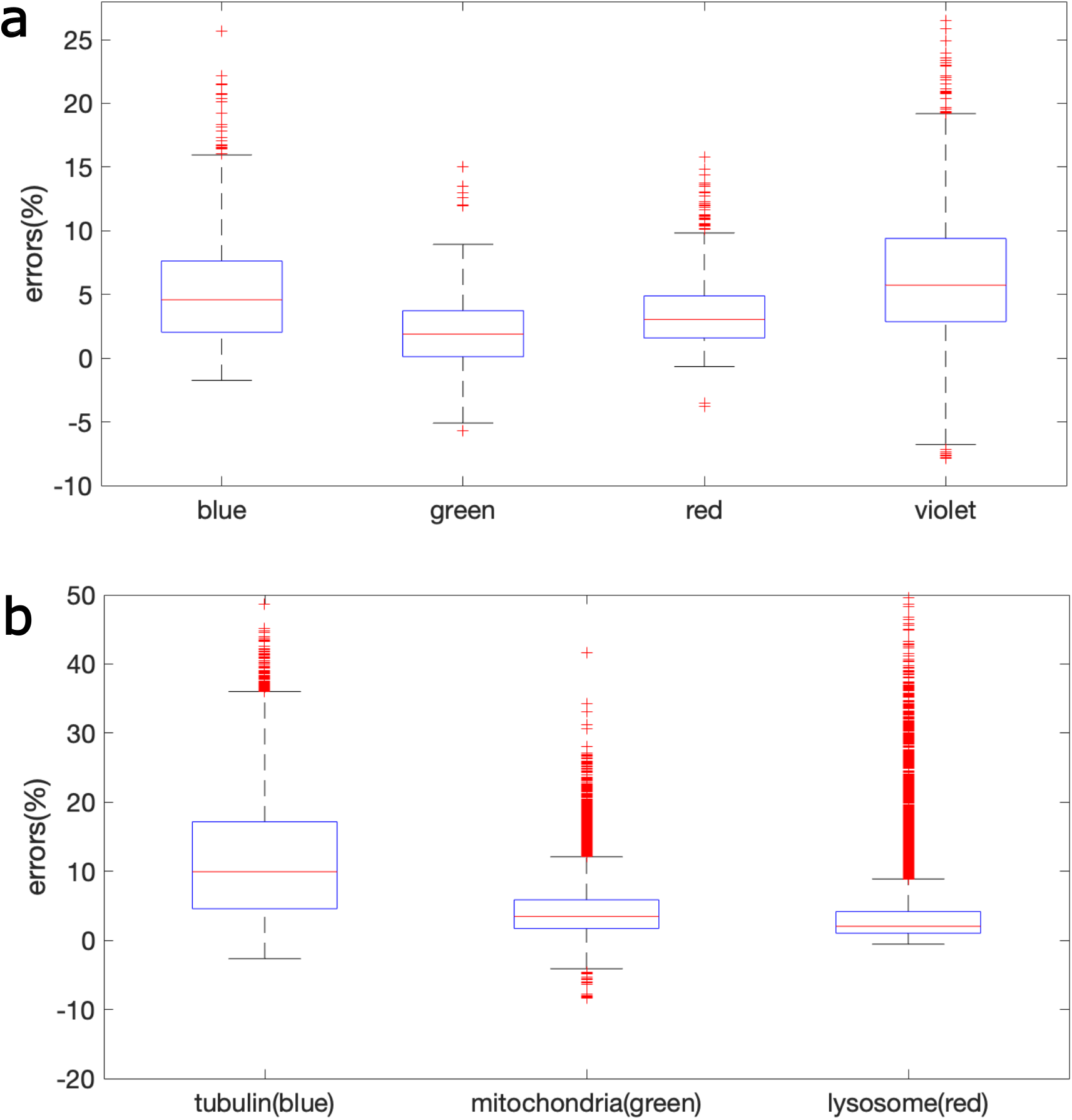
Errors in fluorophore distribution maps given as the scaled difference of ground truth maps and maps obtained using data shown in (a) Fig. 2; and (b) Fig. 4. On each box, the central red line indicate the median, and the bottom and top edges of the box indicate the 25th and 75th percentiles, respectively. The whiskers show the extends to the most extreme not considered outliers, and the outliers are plotted individually using the cross symbol.

Here, we assume an array of L detectors each detecting photons with wavelengths coinciding with a spectral band of Δ*λ* and also the corresponding arrival times. The associated parameters are:

1. Pixels: *i* = 1, …, *I*
2. Species: *m* = 1, …, *M*
3. Spectral bands: *l* = 1, …, *L*
4. Probability of a photon from the *m*th species to be detected in the *l*th spectral band: 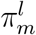
5. Set of 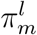 for all *L* spectral bands: ***π***_*m*_
6. Total expected photon count within the *l*th spectral band: 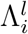
7. fluorophore distribution map associated to the *m*th species within the *i*th pixel: ℐ_*mi*_
8. Lifetime of the *m*th species: *τ*_*m*_ and *η*_*m*_ = 1*/τ*_*m*_
9. Measured photon count from the *m*th species within the *l*th spectral band: 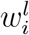
10. The *k*th recorded photon arrival times within the *i*th pixel: 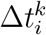

The likelihood over pixel *i* is given as

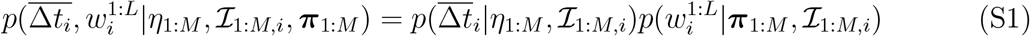

where overbar denotes the set of all possible *k* and^1,2^

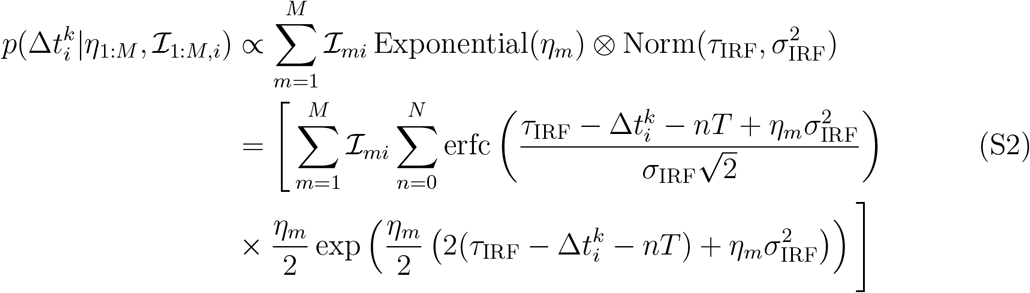

where *n* counts the number of pulses over which the dye remains excited, and

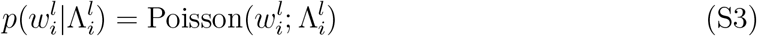

where the total expected photon count within the *i*th pixel and *l*th spectral band is given by

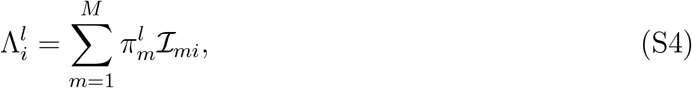

where ℐ_*mi*_ is the total photon count within the *i*th pixel from *m*th species. Moreover

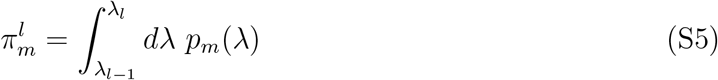

where *p*_*m*_(*λ*) is the emission spectral (wavelength) distribution for *m*th species.

**Fig. S4:**
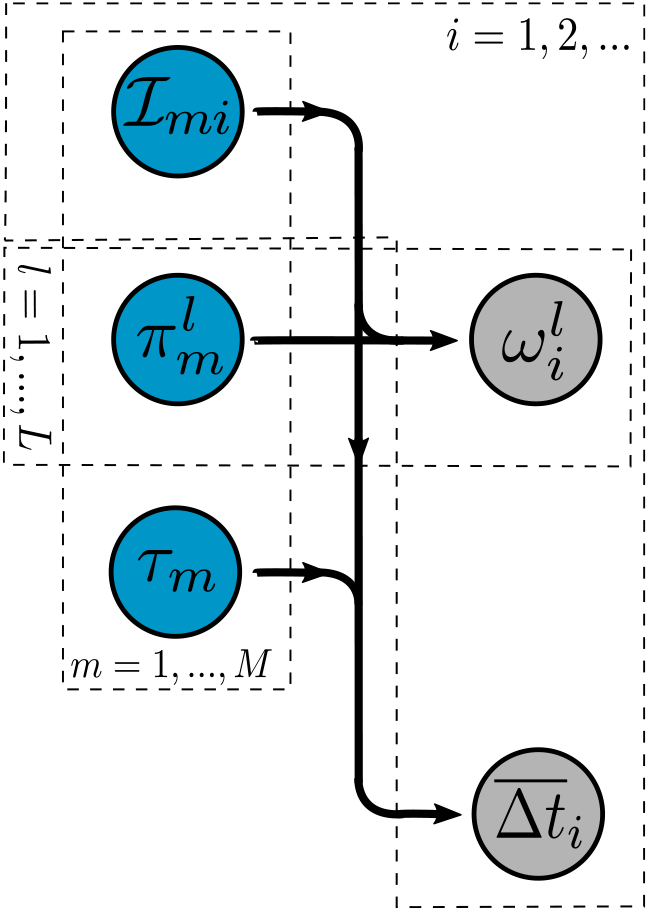
Graphical Model. The gray and blue colors, respectively, denote data and unknowns. The data set includes 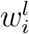 and 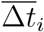 designating the measured photon count from the *i*th pixel and *l*th spectral band, and the set of recorded photon arrival times from the *i*th pixel. 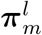 represents the *l*th spectral band of the *m*th species. *τ*_*m*_ and ℐ_*mi*_ are, respectively, the lifetime of the *m*th species and its associated photon count in *i*th pixel. *π*_bg,*i*_ is the fraction of background photons or equivalently the probability of a photon to be originated from background.

From the graphical model, we have the following equations:

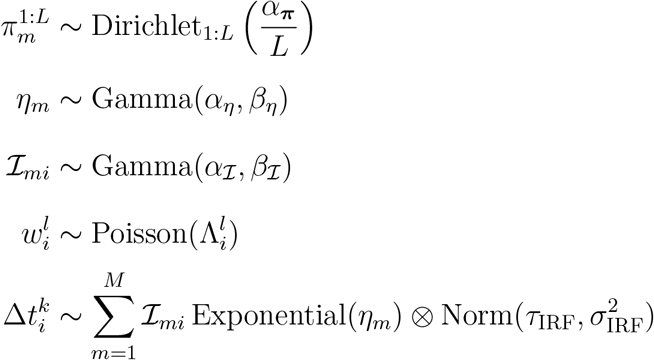

### Inverse Model

#### Sampling *η*_*m*_

The target distribution of *η*_1:*M*_ is given by

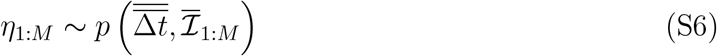

where

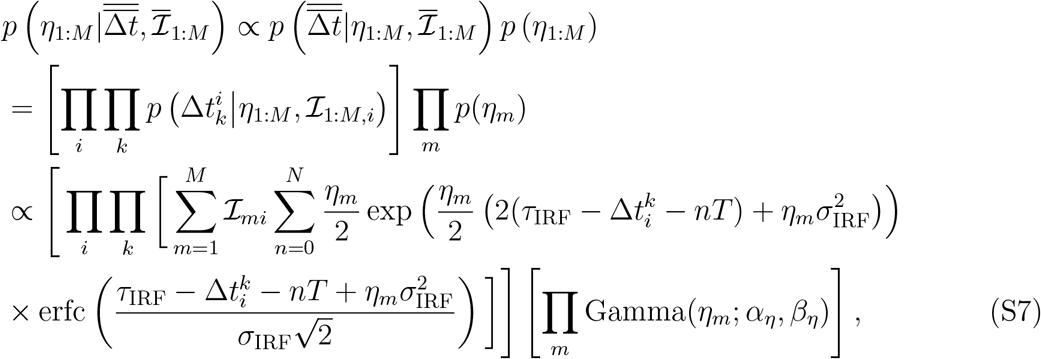

where overbars represents the set of all possible values for *i* or *k*. The above expression does not have a closed form and therefore, Metropolis-Hastings (MH) algorithm is used to sample *η*_1:*M*_. The proposed lifetimes, 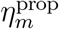, are taken from a Gamma distribution

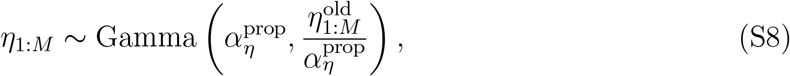

and the MH acceptance ratio is given by

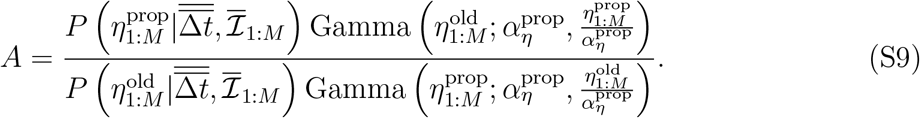

#### Sampling 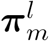

The target distribution of ***π***_1:*M*_ is given by

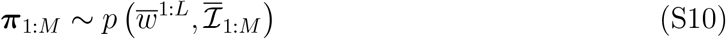

where

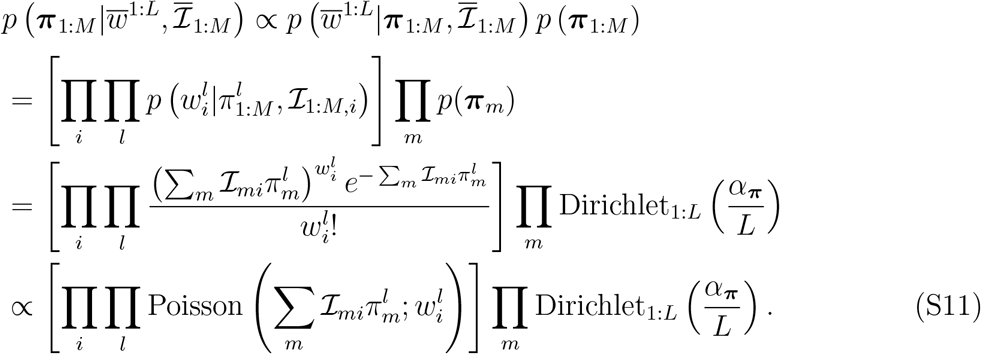

Further, we use a Dirichlet proposal distribution

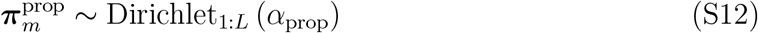

where 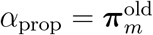, and the acceptance ratio is given as

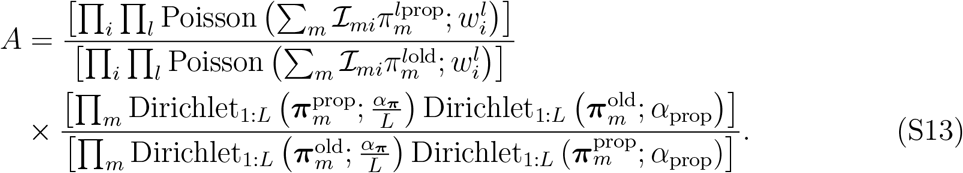

#### Sampling ℐ_*mi*_

The target distribution of ℐ_*mi*_ is given by

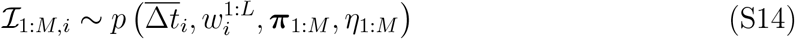

where

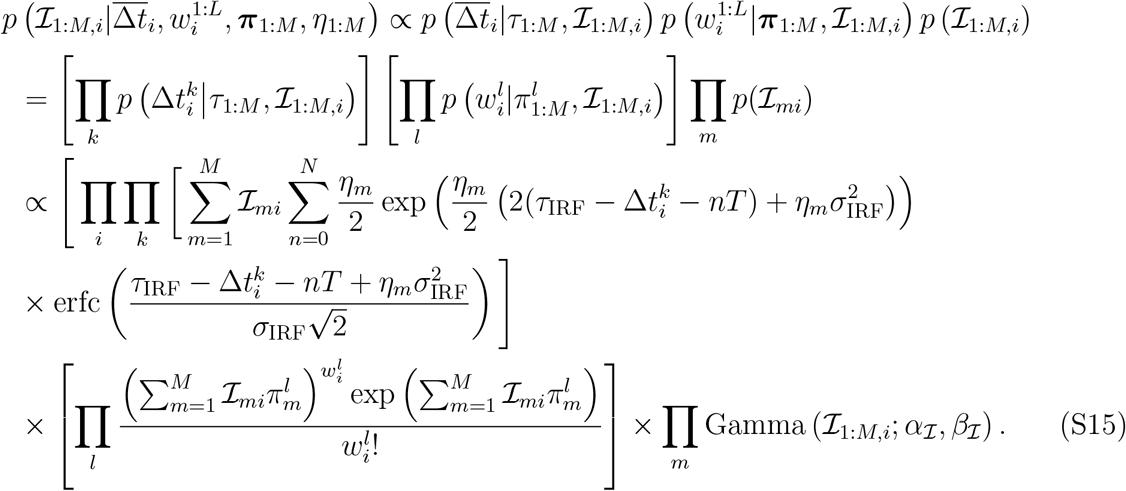

Since the target posterior does not assume an analytic form, we use MH procedure to draw samples. We use a Dirichlet distribution as the proposal distribution

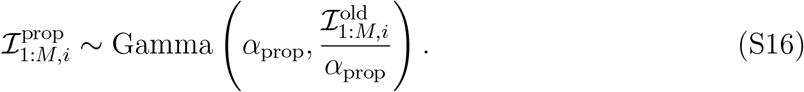

Therefore, the acceptance ratio is

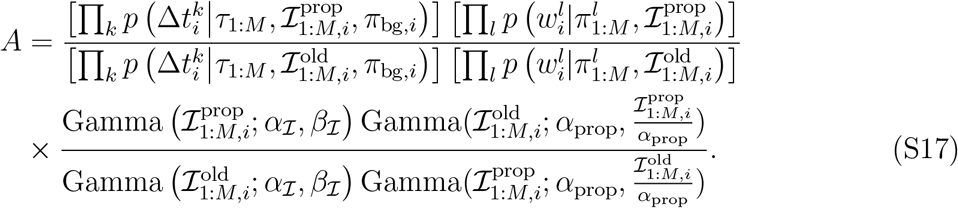

#### Generating Dirichlet random numbers

The Dirichlet random variable is often a vector of *N* random values from the interval [0, 1] that add up to one. Such random vector can be generated as follows:

1. generate *N* random numbers from a Gamma distribution

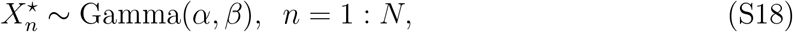
2. divide by the sum of the generated elements to normalize their sum to one

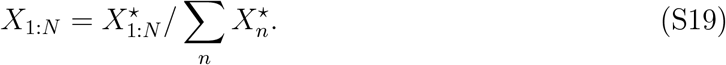

